# Approximate Nearest Neighbor Graph Provides Fast and Efficient Embedding with Applications in Large-scale Biological Data

**DOI:** 10.1101/2024.01.28.577627

**Authors:** Jianshu Zhao, Jean Pierre-Both, Konstantinos T. Konstantinidis

## Abstract

Dimension reduction (or embedding), as a popular way to visualize data, has been a fundamental technique in many applications. Non-linear dimension reduction such as t-SNE and UMAP has been widely used in visualizing single cell RNA sequencing data and metagenomic binning and thus receive many attentions in bioinformatics and computational biology. Here in this paper, we further improve UMAP-like non-linear dimension reduction algorithms by updating the graph-based nearest neighbor search algorithm (e.g. we use Hierarchical Navigable Small World Graph, or HNSW instead of K-graph) and combine several aspects of t-SNE and UMAP to create a new non-linear dimension reduction algorithm. We also provide several additional features including computation of LID (Local Intrinsic Dimension) and hubness, which can reflect structures and properties of the underlying data that strongly affect nearest neighbor search algorithm in traditional UMAP-like algorithms and thus the quality of embeddings. We also combined the improved non-linear dimension reduction algorithm with probabilistic data structures such as MinHash-likes ones (e.g., ProbMinHash et.al.) for large-scale biological sequence data visualization. Our library is called annembed and it was implemented and fully parallelized in Rust. We benchmark it against popular tools mentioned above using standard testing datasets and it showed competitive accuracy. Additionally, we apply our library in three real-world problems: visualizing large-scale microbial genomic database, visualizing single cell RNA sequencing data and metagenomic binning, to showcase the performance, scalability and efficiency of the library when distance computation is expensive or when the number of data points is large (e.g. millions or billions). Annembed can be found here: https://github.com/jean-pierreBoth/annembed

## Introduction

Dimension reduction (DR, or embedding) plays an important role in data science and machine learning, being a fundamental technique for visualization and pre-processing. There are two categories of algorithms for dimension reduction: those seek to preserve pairwise distance for all samples such as PCA and MDS and those seek to only preserve local distance over global distance such as t-SNE (Van der Maaten and Hinton, 2008) and UMAP (McInnes, et al., 2018). The latter is also called non-linear dimension reduction techniques. Recently, non-linear dimension reduction has also been applied to bioinformatics and computational biology such as single cell RNA sequencing analysis (Becht, et al., 2019; Kobak and Berens, 2019) and metagenomic binning (Schmartz, et al., 2022). UMAP was designed to preserve more of the global structure with superior run time performance compared to t-SNE and has no computational restrictions on embedding dimension. For both UMAP and t-SNE, the very first step is to find closest points or neighbors for each point in the dataset. The approximate nearest neighbor algorithm used in UMAP and LargeVis, such as NN-Descent and random projection tree (with neighbor exploring strategy to improve accuracy) are the bounded step for computational complexity (McInnes, et al., 2018; Tang, et al., 2016) and NN-Descent is based on neighborhood propagation to improve accuracy when building K nearest neighbor graph for high dimension dataset. K nearest neighbor graph (K-NNG) was widely used to find nearest neighbors of all elements in a dataset. Popular K nearest neighbor graph construction algorithms including NN- Descent (Dong, et al., 2011), Recursive Lanczos Bisection (Chen, et al., 2009). NN-Descent was used in UMAP to construct K-NNG with an empirical time complexity O (N1.14). This empirical time complexity relies heavily on the properties and distributions of the data. Recently, a new concept called local intrinsic dimension (LID), which was defined as: A data set Ω ∈ ℝ^!^ is said to have local intrinsic dimension (LID) equal to M if its elements lie entirely, without information loss, within a M-dimensional manifold of ℝ^!^, where M < N (Camastra and Staiano, 2016). For example, NN- Descent recall will be very low for dataset with large local intrinsic dimension (LID) such as LID > 20 because it produces large amount of incorrect K nearest neighbors when applied upon high-dimensional data (Dong, et al., 2011; Radovanovic, et al., 2010). Several fast algorithms for building K-NNG has been proposed with time complexity O(d*nt) (1<t<2, normally 1.36 or 1.22 for 2 sub-algorithms) or O(d*N*log(N)) (where d is dimension) and could be further studied for dimension reduction purposes (Chen, et al., 2009; Wang, et al., 2013). The UMAP software implement the NN-Descent algorithm for K-graph building, which is fast for low LID dataset and small K. However, as K or LID increases, the performance of NN-Descent algorithm degrades significantly (Dong, et al., 2011). Another aspect of LID or more generally, the curse of dimensionality, is the hubness concept. Large LID dataset is easier to contain hubs and LID is correlated with hubness. Hubness is defined as the tendency of intrinsically high-dimensional data to contain hubs — points with high in-degrees in the K-NNG, or skewness of the distribution of neighbors of points. NN-Descent on dataset with large LID or hubness is problematic (Radovanovic, et al., 2010), despite some recent efforts to alleviate this problem, such as using a much larger K but reduce the K-NNG to the actual K we need for embedding to improve quality or integrating the information about hubness values into the choice of points to compare with the current approximation of the NN list when build K-NNG (Bratić, et al., 2018). In many real-world datasets, for example, genomic dataset and single cell RNA sequencing dataset obtained by sequencing actual biological samples are distributed highly uneven and biased against well studied organisms or cell types, thus they can be highly clustered and have high LID or contains many hubs.

Finding nearest neighbors based on graph structure, e.g. K-NNG or small world graph has been extensively studied in the past several years (Hajebi, et al., 2011; Wang, et al., 2013) and it turns out that hierarchical small world graph showed high performance and recall in various benchmark studies compare to K-NNG or tree-based NNS search algorithms due to hierarchy and small world property (Aumüller, et al., 2020; Malkov and Yashunin, 2020) . Some modified K-NNG, such as K-NNG + graph diversification and diversified proximity graph (DPG) were shown to have similar performance compare to HNSW, especially for high LID dataset (Lin and Zhao, 2019). As the LID of dataset further increases, the accuracy of HNSW and other graph based methods are compromised if maintaining the same search speed, or the speed will decrease if maintaining the same accuracy/recall, at least for L2 distance (Aumüller and Ceccarello, 2021). Therefore, dimension reduction tools should also consider the LID and hubness of dataset to be embedded for further evaluation of how reliable the nearest neighbor search step can be. There is another UMAP implementations called UWOT and based on ANNOY, which is a random projection forest/tree method for nearest neighbor search instead of NN-Descent (McInnes, et al., 2018). Since ANNOY is a collection of several tree and space partitioning-based algorithms, there is not a clear time complexity analysis. However, benchmark showed that it is much slower than HNSW for high recall for dataset with different LID. Random projection methods such as ANNOY also suffers from high LID and the speed decrease much faster (10X∼100X slower) than HNSW as LID increases given the same accuracy (Aumüller and Ceccarello, 2021). Trimap was also based on ANNOY and was considered faster than UMAP and offers quantitative estimation of how well the global structure was preserved compare to PCA (Amid and Warmuth, 2019). Both HNSW and ANNOY can accelerate nearest neighbor finding in UMAP-like algorithms with time complexity close to O(N*log(N)) (at least for HNSW), where N is the number of database elements. For large dataset, e.g., N >> ∼10^5, N^1.14 (empirical time complexity of K-graph) will be much more expensive than N*log(N), especially when the distance computation is expensive since total time will be number of computations (time complexity) * time for each computation. It has been shown that HNSW, compared to other graph-based nearest neighbor search idea such as NSG (Navigating Spread-out Graph), can alleviate the hubness issue by limiting maximum degree for each point (Zhao, et al., 2023).

It has been shown that t-SNE, combines an attractive force between neighboring pairs of points with a repulsive force between all points (Van der Maaten and Hinton, 2008) . Mathematical analysis has shown that changing the balance between the attractive and the repulsive forces in t-SNE using the exaggeration parameter yields a spectrum of embeddings, which is characterized by a simple trade-off: stronger attraction can better represent continuous manifold structures, while stronger repulsion can better represent discrete cluster structures (Böhm, et al., 2022). UMAP on the other hand, correspond to t-SNE with increased attraction with the same loss function because the negative sampling optimization strategy employed by UMAP strongly lowers the effective repulsion, which leads to more clustered embeddings/structures. However, if we initiate the nearest neighbors from HNSW graphs (instead of initializing from a list of neighbors for each node), it is possible to adopt a different edge and/or node sampling strategy to have similar effects to the attractive and the repulsive forces using the same loss function as in t-SNE/UMAP, by taking into account edge weight distribution and node neighbor density.

Both UMAP and t-SNE were widely applied in single cell RNA sequencing studies since they are much faster than PCA for larger number of cells in single cell RNA sequencing experiments. Visualizing genomic information or metagenomic binning/clustering (e.g., DNA or RNA of cells) using the mentioned dimension reduction technique has several challenges: 1. Genomic/sequence distance such as genomic/sequence identity based on sequences alignment is very expensive to compute via traditional methods such ANI/AAI (for genomes) or alignment identity (for short sequences but not genomes/MAGs); 2. Larger number of genomes in public database (e.g. 10 million for virus database) exacerbate the problem. K-mer hashing based probabilistic data structures (e.g., MinHash) were much faster than traditional ANI/AAI to calculate genomic distance while maintaining ANI/AAI accuracy (Ondov, et al., 2016). Specifically, Jaccard index estimated by MinHash algorithms can be transformed into ANI/AAI/identity via the Mashequation: 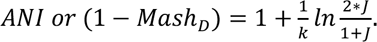. However, they are not combined with UMAP-like fast dimension reduction technique to further accelerate dimension reduction and visualize genomic database. Visualizing microbial genomic database such as GTDB (prokaryotic genome database) (Parks, et al., 2021), IMG/VR (virus species database) (Camargo, et al., 2022) and MyCOcosm (fungal genome database) can help microbiologists and taxonomists quickly check or examine genome affiliation information relative to other genomes according to genomic identity or distance (e.g. ANI or Mash distance) in the database and provide easy-to-catching information about overall database composition, hierarchy and evolutionary space of the target organism (e.g., virus or bacteria). By combining MinHash-like algorithms and HNSW, we created a new data structure to build genomic HNSW graph database (Zhao, et al., 2022) and then apply UMAP-like algorithms to visualizing microbial genomic dataset (aka genomes) to show that when distance computation is very expensive for large number of data points, the nearest neighbor search algorithm becomes an even more limiting step because time complexity of NNS algorithms is directly related to the total number of distance computations and thus total running time.

Here we present a non-linear dimension reduction library called annembed, which utilizes the HNSW for nearest neighbor search for the first step (NNS) and combines embedding techniques from both t-SNE and UMAP. We also add the estimation of hubness and calculation of LID for the database to be embedded compared to the original UMAP. To show applications of the library for biological sequence dataset, MinHash-like algorithms such as ProbMinHash (Ertl, 2020), SuperMinHash (Ertl, 2017), Densified MinHash (Shrivastava, 2017) and SetSketch (Ertl, 2021), are used to estimate Jaccard index for genomic dataset (e.g., microbial genomes such as bacteria and virus database) while Order MinHash (Marçais, et al., 2019) for marker gene dataset , which were then used as input for annembed (that is HNSW were built upon the MinHash-like estimated Jaccard distance) to visualize large scale genomic or marker gene database. The application of annembed library for genomic database is in implemented in a tool called genome embed. Another advantage of annembed is that we use range approximation and approximated SVD (Single Value Decomposition) for dense and/or row compressed matrices to further accelerate UMAP -like algorithms (Halko, et al., 2011), which is widely seen in real world dataset. Annembed is written in pure Rust and can be used as both a stand-alone tool and a library. Annembed is fully parallelized for almost all steps and our benchmark against standard tools showed excellent performance, especially for dataset with billions or more data points, for which current tools were either too slow or failed. Genome embed can now visualize 3 million virus genomes in several hours, much slower if based on UMAP. The idea of visualization genomic database based on MinHash and annembed can be applied to any other sequence database (nucleotide or amino acid, 18S/ITS gene database for example) provided an appropriate distance metric is available to calculate how similar database sequences are to each other (global alignment distance for example).

## Results

### Speed and Visualization Accuracy for Standard Benchmark Datasets

We benchmark annembed with standard dataset MNIST-digits and MNIST-fashion. Annembed performs as good as UMAP (Figure S1 and S2) with similar running time using 24 threads (Table S1). We then test the NNS performance with a large dataset called HIGGS (∼11 million data points, 20 dimensions, generated by Large Hadron Collider) for UMAP (NN-Descent or Annoy for NNS, called UWOT) and annembed, NNS step takes about 18 minutes for annembed while it cannot finish for UMAP (NN-Descent) within 1 hour. For the steps after NNS (e.g. minimize the loss function and produce embeddings), our implementation takes about 43 minutes using 24 threads while UMAP and UWOT takes more than 8 hours (not parallelized) despite the fact that UWOT (ANNOY) NNS step is much faster than UMAP (NN-Descent) because of the ANNOY library: NNS step takes about 22 minutes, similar to that of annembed. However, it has been showing that as K increases (default K is 15 for both UMAP and UWOT), for example K=200 or above, to maintain the same accuracy,, e.g., 95% or higher recall ANNOY is much slower than HNSW according to Aumüller, et al. (2020). Both UMAP and annembed use cross entropy optimization, which is the speed limiting step for UMAP-python and UWOT implementations. However, annembed fully parallelize this step and allow multi-threaded cross entropy optimization, which is about 10X times faster than UMAP for the same step. Despite being fast due to parallelization, memory consumption also increases, we then compared annembed with Trimap (the most memory efficient algorithm for non-linear DR) for embedding the HIGGS dataset, annembed consumed a maximum memory of 58G with 24 threads while Trimap only consumed a maximum memory of 15G (Table S3) but similar running time with UMAP. We also showed that annembed scaled well as the number of threads increases for datasets with millions of data points (Figure S3). The visualization of embeddings for HIGGS following similar pattern for UMAP and annembed (Figure S8 a and b) with UMAP being more compacted.

We then also compare annembed with UMAP using standard single-cell RNA sequencing dataset called PBMC (Peripheral Blood Mononuclear Cells) and C. Elegans. Annembed can clearly separate each cell type of the blood cells from each other and showed consistent visualization with UMAP despite less compacted visualization (Figure S3 (a) and (b)). The C. Elegans dataset also showed consistent result with UMAP, e.g., each cell type and sampling time point are identified as in UMAP (Figure 3 (a) and (b)). Additionally, we can adjust the spatial scale parameter (via --scale) to allow more or less clustered visualization of the embeddings. The default value is used for the above comparison.

### Quality of embedding and estimation of various metrics

We evaluate the quality of embeddings by increasing perplexity (a parameter to balance attention between local and global aspects of the data as in t-SNE). As perplexity quantile increases from 0.05 to 0.99, the quality of embeddings (matched neighbors in embedded space out of 15 true neighbors in the original space) increases from ∼4 to ∼6 for FASHION dataset at quantile 0.5, but did not increase any further as perplexity quantile further increases to 0.99 (Table S3). We also ran the quality evaluation for the GTDB genome database (see below section), the quality of embeddings (matched neighbors in embedded space out of 25 true neighbors in the original space) increased from 12 to 15 as quantile increases from 0.05 to 0.5 but did not further increase as quantile further increases to 0.99 (Table S3). Therefore, annembed can determine the best perplexity to use for different dataset to maximum the quality of embeddings.

LID estimated for MNIST-digits (22.97) and MNIST-fashion (17.5) are very similar to the estimation two related papers (19.6 and 15.3). Hubness estimation in annembed of two standard datasets (2.46 and 1.01 respectively) is also close to the original paper that proposed this concept (Table S2). The hubness of HIGGS dataset is about 1,000 since the number of data points is much larger than MNIST-fashion dataset.

### Visualization large-scale microbial genomic database

We combined MinHash-like algorithms for genomic distance estimation (ANI) with HNSW to obtain nearest neighbor genomes in the database, that is to build HNSW graph using MinHash estimated Jaccard index as a proxy of ANI. For GTDB database (prokaryotic genome database clustered at 95% ANI species threshold), tohnsw (subcommand in GSearch to build HNSW graph) takes about 43 min and 4 minutes for graph building and embedding. For NCBI/RefSeq prokaryotic genome database (∼318K genomes), tohnsw step took about 2.3 hour while embedding step took 13.2 min. Tohnsw step to build HNSW graph database for ∼3 million virus species took about 16.4 hours while embedding step takes about 33 minutes on a 24 threads node. We want to mention that traditional dimension reduction methods such as PCA, NMDS requires calculation all versus all pairwise genomic distance among all database genomes, which will be extremely slow and not practical for real-world genome database (N2 comparisons) because genome comparison is slow even with faster algorithms such as MinHash (Ondov, et al., 2016) for so many comparisons (e.g., 12 million virus genomes, 10^14 comparisons). Hubness estimation is 153.2 at amino acid level for GTDB and 181.2 for NCBI/RefSeq, consistent with the prediction that biological database is highly clustered (MNIST-Fashion has a much smaller hubness estimation, 2∼3, Tables S2, similar total number of data points), whern NN-Descent algorithm performance will degrade significantly.

We then also visualized the embedding result for GTDB, which is a taxonomically well described (labeled) genome species database according to ANI/AAI distance. It is clear that a majority of the genomes were well visualized by annembed according to the taxonomy affiliation of each genome, e.g., genomes within the same phylum affiliation sit in the same cluster in the 2 dimension annembed plot (Figure 2). However, we also observed that some of the genomes, for example, *Firmicutes* (red) and *Desulfobacterota* (orange) genomes sit in the cluster of *Proteobacteria* (green), which indicates that those genomes are more similar to Proteobacteria genomes than their current affiliation in term of their AAI. We then extract those *Firmicutes* and *Desulfobacterota* genomes (3 genomes as an example) and calculate universal gene AAI (accurate for phylum level genomic distance/identity) between them and the most similar genome in *Proteobacteria.* We found that those genomes have larger universal gene AAI values to the most similar genome in *Proteobacteria* than to the most similar genome in their original affiliation (57.3%, 49.5% and 54.9% versus 50.4%, 42.0% and 47.2%). We then also visualized detailed embedding results at a lower taxonomic rank (e.g., class and order level) and found that annembed can also clearly differentiate between different classes and orders in phylum *Proteobacteria* (Figure S4). Therefore, the annembed results can be used to quickly determine or check mislabeled taxonomy information in public databases. We then also visualize the NCBI/RefSeq prokaryotic genome database and closely related genomes sit within the same cluster at phylum level (Figure S5), consistent with the GTDB database visualization. We then also visualize the virus genome database, with closely related virus genomes sit in the same cluster (Figure S6).

### Visualizing Large-scale marker gene databases for prokaryotes

We combined Order MinHash with annembed, a LSH to approximate edit distance (an alignment-based distance), which can be used to approximate alignment identity (Marçais, et al., 2019), to visualize 16S ribosomal RNA database (e.g., SILVA, RDP) (average ∼1,500 bp). Since Order Minhash is a special case for the weighted MinHash (weight is the position of the kmer in a sequence, but not abundance of the kmer), we applied the same computational techniques as in ProbMinHash to accelerate hashing computation. We can now run annembed with Order Minhash as underlying sequence distance estimation for 1.6 million 16S RNA sequences in SILVA in less than 5 minutes with 24 threads (NNS step 33 minutes). We did not see a clear separation by their corresponding label at phylum or class level (Figure S9 and Figure S10), indicating that many of the taxonomy annotations in SILVA are wrong, consistent with the pioneering analysis of (Edgar, 2018), which concluded that 20% of the taxonomy annotations in SILVA are wrong by looking at the guiding tree. We also ran the same step for RDP v18 16S rRNA database as a comparison. We see a clear separation by taxonomic annotation at phylum, class and even genus level (Figure 11 and Figure 12). Those sequences in RDP are taxonomically identified based on isolate and type strain sequences and/ or environmental sequences predicted by the RDP Naive Bayesian Classifier (Wang, et al., 2007), the nomenclature of which is based on Bergey’s Manual. RDP Naive Bayesian Classifier and the trained database are thought to be one of the most accurate sequence classification system (Edgar, 2016).

### Metagenomic binning via embedding contig kmer composition

t-SNE and UMAP has been applied to help manually binning/clustering metagenomic contigs to obtain MAGs (e.g., plot kmer coverage versus kmer composition) because the dimension reduction based on PCA requires all versus all distance computation, which is computationally expensive for thousands of contigs. Here, we replace t-SNE and UMAP in the metagenomic binning software mmgenome (Karst, et al., 2016) with annembed (LSH for Euclidean distance as contig distance estimation method). Annembed with LSH reduced the computation time to calculate contig profile differences (Euclidean distance based on kmer composition) and dimension reduction by 3-5 times for a medium size metagenomic assembly (10,000 contigs, average ∼8,000 bp) (Tables S4). For even larger metagenomic assemblies (e.g., millions of contigs), annembed will be even faster than t-SNE or UMAP, which can accelerate metagenomic analysis. We obtain similar results with annembed compared with UMAP when manually check the binned MAGs for most of the MAGs/bins (Figure S13). However, we noticed that there are several MAGs, annembed cannot tell whether those contigs are from separated clusters or MAGs but UMAP can tell. We suspect this is because the loss function in UMAP put more weight on the repulsive force, leading to more compact visualization despite the actual distance or similarity for those contigs are not preserved in UMAP, see more discussion on the loss function section.

## Discussion

In this study, we improve UMAP-like algorithms by applying fast and efficient graph-based neighbor search algorithm and providing additional parameter estimations for large scale non-linear dimension reduction tasks. Despite the fact that replacing K-graph with HNSW will greatly accelerate large-scale nearest neighbor finding, HNSW requires that the distance used must be metric (to maintain neighborhood diversity and thus high accuracy), which is not a requirement for NN-Descent algorithm (Dong, et al., 2011). However, the weakness of NN-Descent algorithm is that for high LID dataset, nearest neighbor accuracy and speed decrease significantly and it will be much slower the large datasets (e.g., millions). Importantly, metric distances are more widely applied (e.g. L1 and L2, angular and hamming distance) than non-metric distances in real-world machine learning tasks (Li and Li, 2022). We have shown that in real world dataset with millions of data points, annembed library is at least 8∼10 times faster while maintaining similar visualization accuracy due to both HNSW and improved/parallelized embedding algorithm. Annembed will be even faster as database size keeps growing becase of the O(N*log(N)) HNSW graph database build step. The LID and hubness estimation we provided offer more information about the structure and distribution of the data, especially high dimensional ones, thus help to perform better nearest neighbor search. Those two features were not included by either the original NN-Descent paper or UMAP paper, but a bunch of papers published recently have shown that they affect the true neighbor found for each data point (Bratić, et al., 2018; Radovanovic, et al., 2010) and also affect the performance of NNS finding (Aumüller and Ceccarello, 2021), a limiting step for UMAP-like or t-SNE-like algorithms. SpaceMAP considered LID when performing the space expansion to account for a geometrical distortion between high-dimension space and embedded space (Zu and Tao, 2022). The MLE estimation of LID is widely applied in many cases (Camastra and Staiano, 2016; Levina and Bickel, 2004; Zu and Tao, 2022). However, it requires at least 20 neighbors to be accurate. This is not a problem for testing datasets, but it is sometimes a big problem for biological/genomic datasets because there are not so many accurate genomic distance estimations for distantly related genomes or new genomes: all genomic distance estimation algorithms for nt such as alignment-based ANI, fastANI or Mash will lose accuracy below ∼70% ANI thus not performed when there are such distantly related genomes to other genomes in database (Parks, et al., 2021). We can use the amino acid level genomic distance (e.g., AAI) to alleviate this problem or apply newly invented LID estimation algorithm that does not require at least 20 neighbors (Amsaleg, et al., 2019).

The speed of NNS finding is exceptionally important for real world applications like genomes and other format of data points, instead of just vectors (like those standard benchmark datasets) because total running time is related to the number of comparisons for the entire database and single pair comparison. With a O(N*log(N)) complexity, HNSW is orders of magnitude faster than NN-Descent for large database and it is fully parallelized via Rust, a difficult parallel task for other languages like C/C++ due to clear data race when multiple threads work on the same graph data structure. Thread safe languages like Rust and Java are more convenient to parallelize such tasks. Also, for large dataset, memory requirement for NNS finding is a key limiting step for both NN-Descent and Annoy. Here we implemented memory map in the hnswlib-rs library for running datasets with billions of data points without memory limit problems. More importantly, we provide a trait in the hnswlib-rs library to allow users to implement their own distance. This provided an opportunity to combined MinHash like probabilistic data structures with HNSW since set similarity, for example Jaccard index, can be estimated in a sub-linear running time using the mentioned probabilistic data structures. We provide several MinHash-like implementations as part of the distance estimation step, aiming at accuracy or space-efficiency purposes. The application of annembed to visualization of genomic database based on the idea mentioned above, can be very useful to check the label accuracy of taxonomically annotated databases. For example, we found that many genomes labeled as phylum *Desulfobacterota* are actually *alpha-Proteobacteria* (a class of phylum *Proteobacteria*) (Figure 3). Also, very high hubness for genomic datasets (GTDB or RefSeq) indicates either discontinuous evolutionary space (e.g, frequent genetic exchange within clusters) (Jain, et al., 2018; Koonin and Wolf, 2008) or biased sequencing/sampling efforts towards existing genomic sequences (Murray, et al., 2021) at both species level or high taxonomic rank. More broadly, the mentioned MinHash-like algorithms combined with HNSW for annembed, can also work for text/document files or websites, but not just only genomes, providing an opportunity to run dimension reduction and visualization for large-scale real-world datasets, with the potential speedup many industrial applications.

We also want to mention that in addition to Jaccard index and Edit distance (string matching), Euclidean distance, Hamming distance and Angular distance can all be computed via hashing- like algorithms (Argerich and Golmar, 2017; Datar, et al., 2004; Pacuk, et al., 2016; Pagh, 2016), making the idea of combining them with HNSW even more attractive when we want to run UMAP-like algorithms for those distance metrics for various real-world datasets (e.g., strings, vectors and text/document). We provide an example to use order MinHash to approximate Edit distance for DNA sequences and visualize them via annembed. This application helps to identify mislabeled taxonomic information in widely used reference sequence databases. However, when distance of interest is not a metric distance, HNSW will suffer and UMAP based on NN-Descent will be a better option (NN-Descent works for non-metric distance) even though there are recent efforts that generalize HNSW to non-metric distances (Tan, et al., 2021).

We combined embedding strategies from both t-SNE (e.g., node neighbor density sampling or a probability normalization constraint with respect to neighbors) and UMAP (e.g., cross entropy optimization) but initiate from the HNSW graph (a kNN graph like structure can be extracted from the HNSW graph with constant time), which provides convenience to control the balance between attractive and repulsive force during edge sampling step as in UMAP paper and the same loss function with UMAP was used for cross entropy optimization (McInnes, et al., 2018). It was mentioned recently that UMAP implementation (umap-python and UWOT) did not use the loss function in the paper but a different one, which increase the repulsion on all pairs of points, leading to more compacted visualization but lose high-dimension similarity (Damrich and Hamprecht, 2021), consistent with our observation. But it might be useful to change the optimization procedure of UMAP so as to decrease the repulsion per edge only on the non-kNN graph edges and not on the kNN graph edges as proposed in the new loss function (Damrich and Hamprecht, 2021). We will explore the new loss function proposed to correct the repulsion weight for edges in the shared kNN graph. Both edge sampling and cross entropy optimization strategies we implemented are parallelized and can be much faster than available implementations with more CPU threads, as shown in HIGGS dataset (Figure S3), making the library a good fit for large-scale datasets.

Despite the fact that we provide an implementation to estimate the quality of the embeddings by looking at how many neighbors of each point in original space are preserved in the embedded space locally (similar to the KNN accuracy index to evaluate local structure as in SpaceMAP (Zu and Tao, 2022) ), there isn’t an explicit index to evaluate how well overall annembed preserved the global structure compare to the original space like TriMap and SpaceMAP, which implemented an index to calculate how well the global structure is preserved by using PCA as a standard (Amid and Warmuth, 2019). We will further explore this idea to also provide a similar index to calculate how well the global structure is preserved. TriMap is more memory efficient via triplets constraint (Amid and Warmuth, 2019) but still it is not parallelized, and our results showed that it is 10 times slower for the HIGGS dataset than annembed despite a little bit more memory is required. Even more interesting UMAP-like algorithms such as SpaceMAP, which considered not only a restricted k−nearest neighborhood (part of the manifold, thus ignoring the hierarchical structure) by matching the “capacity” of high- and low-dimensional/embedded space via analytical transformation of distances adjusted by LID (Zu and Tao, 2022), are also not parallelized and thus slow for large datasets. It turns out that for disjoint manifold (MNIST fashion/digits), UMAP, t-SNE and annembed worked well but for continuous manifold (ImageNet (Deng, et al., 2009)), SpaceMAP will be better options. We are not sure whether the evolutionary space is disjoint manifold or continuous manifold given the little sequencing efforts in a huge microbial evolutionary space.

New metagenomic binning algorithms such as Rosella (Newell, 2023), BinArena (Pavia, et al., 2023), mmgenomes (Karst, et al., 2016) relies on UMAP or t-SNE for visualization of contigs after obtaining the composition of contigs (e.g., Euclidean distance based on kmer composition of contigs) to avoid all versus all distance computation. Then contigs will be manually binned via human intuition or clustering algorithms. As the metagenomic sequencing capabilities for a give sample keeps growing, it is possible to have millions of contigs assembled and to be binned. Annembed can be much faster in those cases. However, the distance among contigs in the embedded space of UMAP or t-SNE is not the actual genomic distance in the original Euclidean space consisting of kmers. It is the same with annembed because the global distance is not well preserved. As discussed above, consider non-KNN graph edges in loss function (Damrich and Hamprecht, 2021) or compensate for underlying distortion of distance metric between original space and embedded space as mentioned in SpaceMAP.

It has to be noted, however, HNSW graph building is still not so fast for large datasets due to O(N*(log(n))) despite it is one of the fastest nearest graph building algorithms. Recently, a new algorithm combining LSH with HNSW or approximate proximity graph (APG) structure to further accelerate graph database building step from O(N*log(N)) to O(N*c), where c is a constant independent of N. LSH-APG builds an APG via consecutively inserting points based on their nearest neighbor relationship with an efficient and accurate LSH-based search strategy (Zhao, et al., 2023). A high-quality entry point selection technique and an LSH-based pruning condition are developed to accelerate index construction and query processing by reducing the number of points to be accessed during the search for each query. Annembed could be further accelerated by adopting this idea when building HNSW graph to break the O(N*log(N)) run time limit.

We believe that annembed library will greatly accelerate large scale non-linear dimension reduction tasks (e.g., millions or even billions), where the original one and other methods were limited by the nearest neighbor search step and single threaded implementation following after. It can also help visualizing large-scale genomic databases or single cell RNA sequencing studies and advance scientific discoveries related to cancer and other diseases.

## Methods & Materials

Overall, our implementation is a mixture of the various embedding algorithms such as UMAP and t-SNE. First, the graph is initialized by the HNSW algorithm (Figure 1a), which provides for free, sub-sampling in the data to be embedded by considering only less densely occupied layers (the upper layers). This corresponds generally to a subsampling of 2%-4% but can give a guarantee as the distance between points leaved out the sampling and its nearest sampled neighbor are known. The HNSW structure thus enables also an iterative/hierarchical initialization of the embedding by considering an increasing number of layers (until layer 0). The preliminary graph built for the embedding uses an exponential function of distances to neighbor nodes (as in UMAP) but keeps a probability normalization constraint with respect to neighbors (as in t-SNE) (Figure 1b). It is then possible to modulate the initial edge weight by considering a power of the distance function to neighbors or increase or decrease the impact of the local density of points around each node (similar to the repulsive force). We initialized embedded space by diffusion map (Coifman, et al., 2005) instead of Laplacian Eigenmap as in UMAP, the former can be considered as a generalization of the latter but the order of top eigenvector is reversed. To minimize divergence between embedded pace and initial distribution probability, we also used a cross entropy optimization of this initial layout but consider the initial local density estimates of each point in the embedded space when computing the Cauchy weight of an embedded edge as in UMAP (Figure 1b). The corresponding “perplexity” distribution (the same as in t-SNE, a parameter to balance attention between local and global aspects of the data as in t-SNE) is estimated on the fly. We provide a tentative assessment of the continuity/quality of the embedding by varying the perplexity to help selecting among varying results between runs for a given dataset. Quality of embedding was defined as the correct neighbors of node in the embedded space when comparing with the original data. During this process, LID and hubness were also estimated based on the HNSW graph (Figure 1c and d). See Figure 1 for the overall description of annembed.

**Figure 1.**
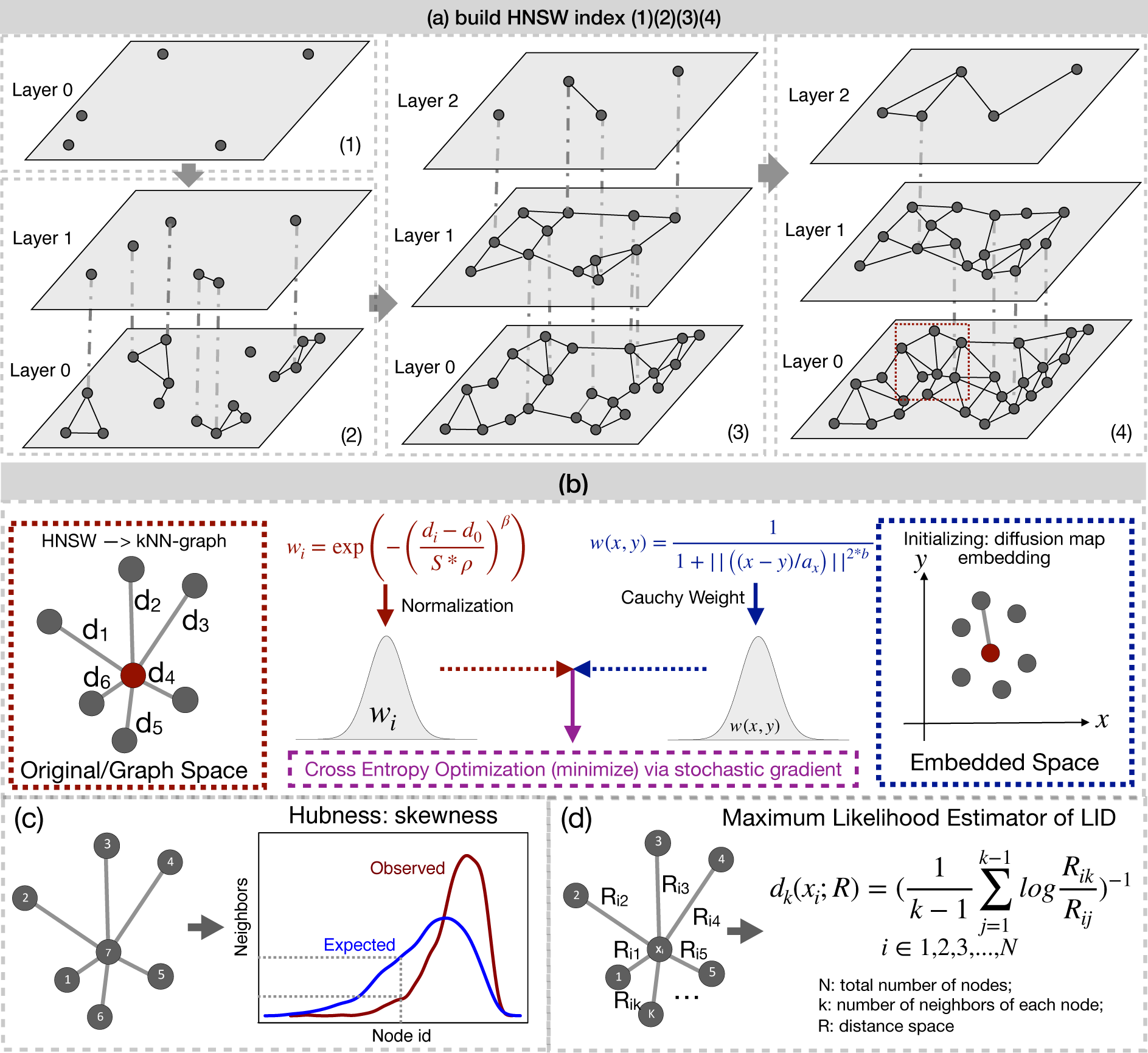
Overall description of annembed key steps/algorithms and functionalities. **(a)** Build HNSW graph from scratch by gradually adding points in the database in a recursive way with random initialization. When maximumly allowed number of neighbors (M) for each existing point was reached in the graph, a representative will be chosen as new point in new layers (above) by collapsing the neighbors. Finding neighbors for a newly added point is to search the graph but the new one will be inserted into the graph after required number of neighbors were found and the graph will be updated (newly added point could also be neighbors of other existing points); **(b)** UMAP-like embedding algorithm by combining t-SNE (edge sampling in left panel) and UMAP (cross entropy optimization). For edge sampling, β is set to 1 to so that we have exponential weights similar to Umap. *S* is to modulate ρ and is set to 0.5 to while ρ is the spacial scale factor and is also set to 1. After normalization the weight will be probability distribution; For initial embedding, to define the weight of the edge, we initialize *b* as 1 and coefficient *a*_*x*_ is a coefficient related to scale coefficient in the original space around x since scales are smooth as they are computed as mean of distances around x: for each point x we have its k neighbours yi [i=1,2,3…k], for each of its neighbours yi we get its first neighbour distance di. We then average all these distances, that give us an idea of distance around a point, a scalex, 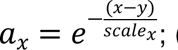 **(c)** Hubness estimation by evaluating the skewness of neighbor distributions of observed neighbors of each node and the expected ones. **(d)** LID estimation via the maximum likelihood method. We use Euclidean distance (metric) for R by default and it can be changed according to user specific case (e.g., Jaccard distance, a metrric, can be used for genomes).

**Figure 2.**
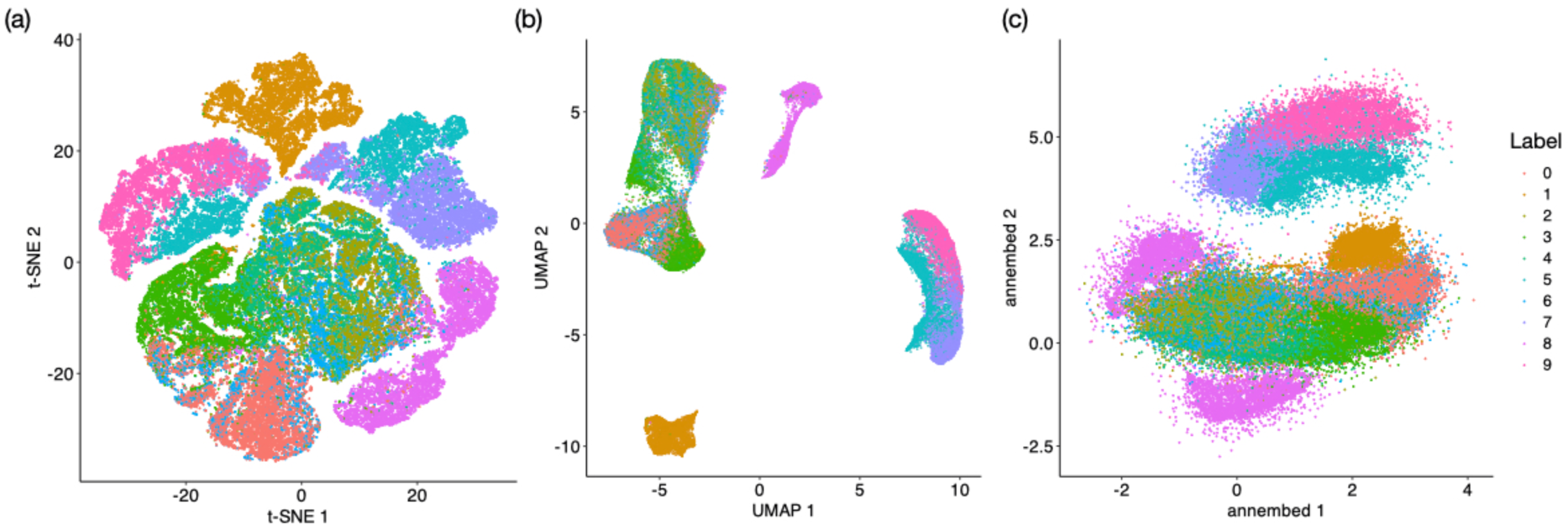
Dimension reduction for t-SNE (a), UMAP (b) and annembed (c) respectively for MNIST- fashion dataset. Color legend indicates different labels. For t-SNE, UMAP and annembed, k=15 (number of neighbors) was used.

**Figure 3.**
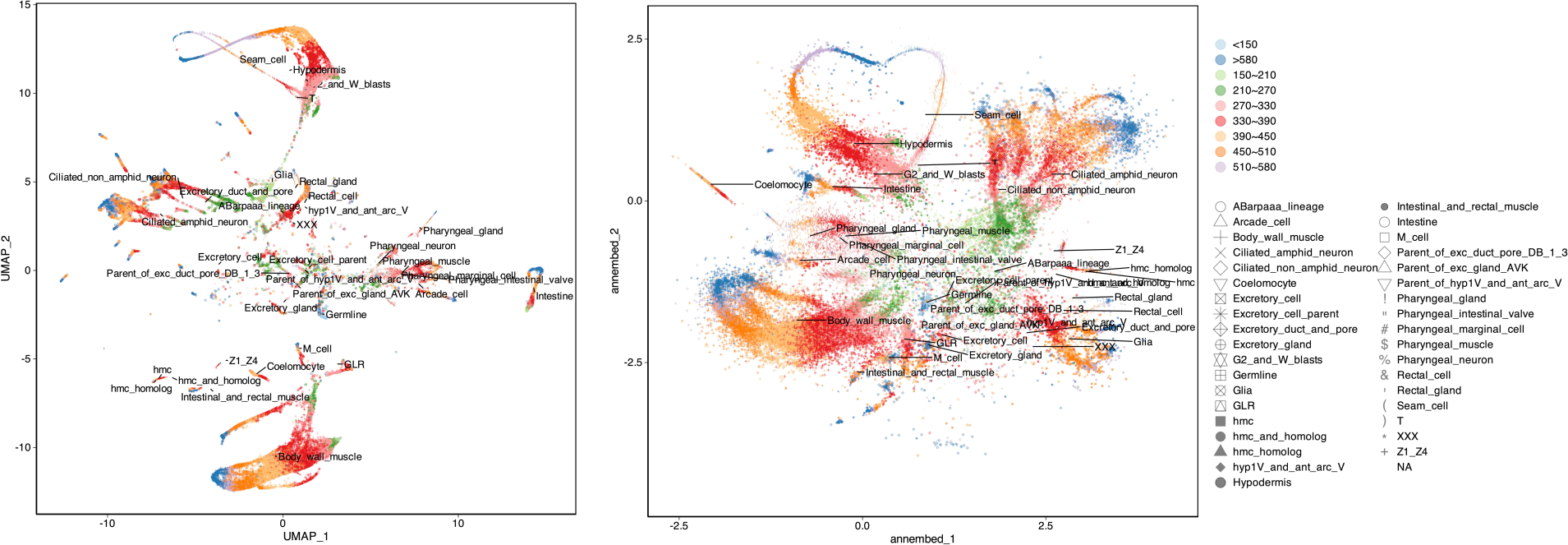
UMAP and annembed visualization for C. Elegans single cell RNA sequencing dataset. Color indicates time (h) since experiment started while shape indicates cell types. Major cell types are also labeled in the plot, anchored by the centroid of each cell type. High resolution figures can be found in the supplementary materials.

**Figure 4.**
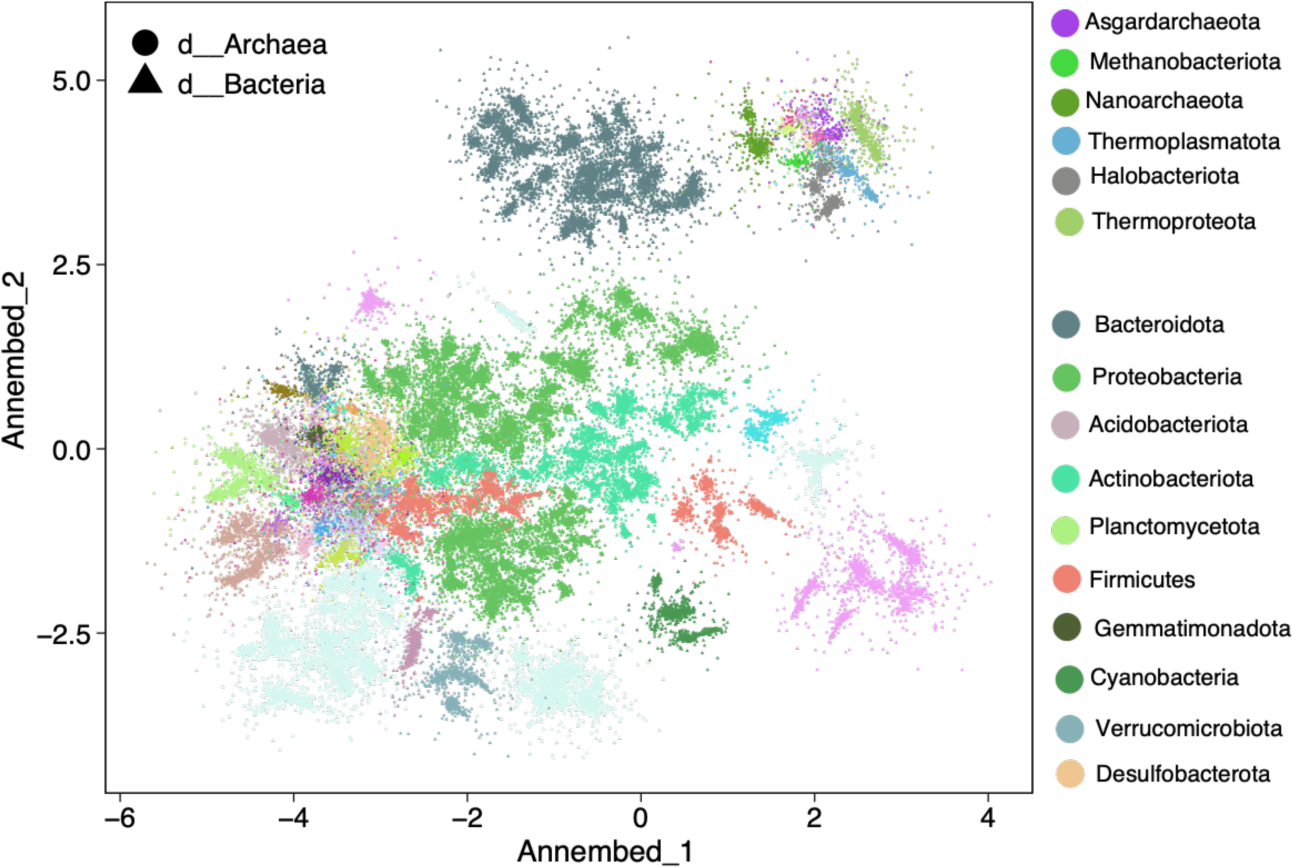
Annembed plot of GTDB v207 (65,703 genomes) using ProbMinHash as genomic distance metric. Different shapes indicate bacteria or archaea kingdom. Major phylums in both bacteria and archaea kingdom are colored accordingly. ProbMinHash distance was based on predicted gene amino acid sequences. We cannot run on nucleotide level genomic distance because a pre-clustered database at 95% ANI is too sparse, many of the genomes do not have reliable genomic distance estimation at nucleotide level to meet the minimum requirement of 15 nearest neighbor genomes in UMAP and annembed.

### Hierarchical Navigable Small World Graph (HNSW)

We use HNSW instead of K-graph to find nearest neighbors for each data point in the dataset to be embedded. Specifically, HNSW incrementally builds a multi-layer structure consisting of a hierarchical set of proximity graphs (layers) for nested subsets of the stored elements, which maintains the small world property (Figure 1 (a)). Then, through smart neighbor selection heuristics, inserting and searching the query elements in the multi-layer proximity graphs can be very fast (due to small world property for each layer and hierarchical structure) while preserving high accuracy/recall. Inserting new data into existing graph is essentially a search process but all neighbor list in the graph will need to be updated once the inserting is completed. We reimplemented the hnswlib library in Rust and benchmark it against standard datasets. Note that HNSW require metric distance as input because the heuristics in maintaining neighborhood diversity requires metric distance. Build the graph takes O(N*log(N)) time. We then extract K neighbors of each point/node in the graph database for embedding as mentioned above. Note that building HNSW is the same to search a new element against graph database except that for building, all elements in database need to be searched in a recursively way, during which database graph needs to be updated after search is done for each database element. The hnswlib-rs can be found here: https://github.com/jean-pierreBoth/hnswlib-rs

### Embedding and quality of the embedding

Embedding was done by the following steps: 1. Initialize from HNSW graph, an exponential function of distances to neighbor nodes for all nodes was calculated but keeps a probability normalization constraint with respect to node’s neighbors (Figure 1(b) left pannel); 2. Adjust the initial edge weight by considering a power of the distance function to neighbors (increase or decrease local density of points around each node) for the embedded space (Figure 1(b) right panel); 3. Minimize divergence between embedded and initial distribution probability, then perform cross entropy optimization of this initial layout but consider the initial local density estimates of each point when computing the Cauchy weight of an embedded edge (Figure 1(b)); 4. Estimate the corresponding “perplexity” distribution. 5. Varying perplexity and evaluate the quality of embeddings as follows. We use the same loss function for divergence minimization or cross entropy optimization as in UMAP paper (McInnes, et al., 2018) (see Supplementary Methods).

To quantify the quality of the embedding, annembed tries to assess how well neighborhood of points in original and neighborhood of points in embedded space may match, also called KNN accuracy. In each neighborhood of a point, taken as center in the initial space, we search the point that has minimal distance to the center of the corresponding neighborhood in embedded space. The quantiles on this distance are then dumped. We also count the number of points for which the distance is in the radius of the neighborhood (of size number of neighbors in HNSW graph) in embedded space and for neighborhood that have a match, the mean number of matches. This will then provide a continuity/quality of the embedding.

### Hubness

LID is directly related to another important concept called hubness, defined as the tendency of intrinsically high-dimensional data to contain hubs — points with high in-degrees compared to the average in-degrees in the K-NNG (Radovanovic, et al., 2010). It has been shown that those two are correlated. NN-Descent (implemented in UMAP), as the key algorithm to build K-graph, performs very bad for such dataset with hubness, that is a skewed distribution of point neighbors when compared to an expectation (some might have much more neighbors than others) (Dong, et al., 2011). We estimated the skewness of point neighbors of the dataset by comparing the neighbors we actually observed with the expected neighbors (e.g. average number of neighbors) according to the original paper in our library (Radovanovic, et al., 2010) (Figure 1(c)): At the very beginning, we initialize the hubness values of each data set point to zero. Then, during algorithm execution (K-NN was extracted from HNSW), we increase the hubness value of a given point by one if that point is added to the nearest neighbor list of some other point, and analogously, we decrease the hubness value by one if the point is removed from some nearest neighbor list.

### Local Intrinsic dimension (LID)

To estimate the local Intrinsic dimension (LID) of a dataset that users want to embed, we implemented the maximum likelihood estimation (MLE) (Aumüller and Ceccarello, 2021; Levina and Bickel, 2004) after comparing it with other estimations such as Methods of Moments (MoM), Probability-Weighted Moments (PWM) and Regularly Varying Functions (RVF). The MLE estimation has a significantly smaller standard deviation compare to other methods and converge faster to mean than other methods as the number of sample increases (Amsaleg, et al., 2015). Note that by default the MLE estimation needs more than 20 neighbors around each point to have reliable estimation. There is a new algorithm called tight localities that can have more reliable estimations even for neighbors smaller than 20, we plan to also implement it in next release (Amsaleg, et al., 2019). Specifically, the MLE method to estimate the ID was based on constant density assumption in a small neighborhood and the Poisson process to model the random sampling in this neighborhood (Levina and Bickel, 2004). The MLE method provides a way to estimate ID for point xi in its k−neighborhood (k >= 20). Let R be the distance metric (e.g. Euclidean) and Rij ∈ R be the distance between point xi and xj under this metric, the maximum likelihood estimation (MLE) of the ID around point xi, with the distance metric of R, is computed as: 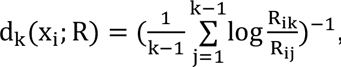, where the summation is over the k-nearest neighbors of point x.

We note that d_k_(x; R) is point-specific, dependent on k and the distance metric R (Figure 1(d)). LID therefore uniquely characterizes the sub-manifolds around x. We then average this value across all point in the database.

### MinHash-like algorithms for genomic distance estimation to approximate ANI/AAI for genomic visualization

MASH is a fast and efficient software to compute genomic distance via MinHash algorithm and correlates well with the golden standard of genomic distance measurement ANI after transformation based on evolution model 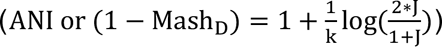 (Ondov, et al., 2016). However, MinHash does not take into account the kmer abundance (or multiplicity) and also total kmer count of a given genome, which affect the estimation of real genomic distance of genomes (Koslicki and Zabeti, 2019; Rowe, et al., 2019). To consider multiplicity of k-mers in the k-mer set of genomes, traditional hashing algorithms will not be a good choice since they all assume unique set element (k-mer). New MinHash algorithms were designed for weighted set to solve the above-mentioned problem (Christiani, 2020; Ertl, 2018; Ioffe, 2010; Wu, et al., 2020).

Still, those weighted MinHash algorithms do not solve the problem of different set size in biasing estimation of weighted Jaccard index (Koslicki and Zabeti, 2019; Rowe, et al., 2019). Recently, ProbMinHash was proposed to take into account both weighted set (k-mer multiplicity) and total set size (total k-mer count) (Ertl, 2020). Accordingly, new Locality Sensitive hashing algorithms (P-MinHash) for weighted set and different set size was proposed to estimate weighted and normalized (to account for set size difference) Jaccard like index Jp (Moulton and Jiang, 2018; Yang, et al., 2019) and ProbMinHash was built upon those new hashing algorithms with further computational optimization. More importantly, it is a proper metric distance and can be used as input to HNSW. We also implemented a more accurate MinHash-like algorithm called SuperMinHash (also a locality sensitive hashing) for simple Jaccard index estimation (Ertl, 2017) but it is slower. SetSketch, a MinHash-like, but a combination of MinHash and HyperLogLog, is also implemented for its space efficiency (e.g., require much less memory and disk space to store the sketched genomic information)(Ertl, 2021). Another much faster than but as accurate as traditional MinHash (as in Mash) called One Permutation MinHash with Optimal Densification (Densified Minhash) is also implemented (Shrivastava, 2017). We have shown that ProbMinHash/SuperMinHash/SetSketch/Densified MinHash distance correlates very well with ANI and AAI for 80-100% ANI and 55%-95% AAI after transformation (Zhao, et al., 2022). To approximate sequence alignment (contiguous sequence like 16S rRNA) identity via MinHash-like algorithms, the Order MinHash (Marçais, et al., 2019) was implemented, which allows fast and accurate computation of edit distance based on kmers set distance but not alignment. Order MinHash is very similar to that of ProbMinHash except that the weight is the position of the kmer and that normalization by total kmer count is not necessary. All those implementations are available in the probminhash library (https://github.com/jean-pierreBoth/probminhash).

### Linear Algebra Backend

The annembed package involves basic vector and matrix operations for UMAP algorithm (e.g., minimize a target function), which is computationally intensive for large dataset and thus we use high performance linear algebra libraries such as BLAS. For Intel processors, we use Intel Math Kernel Library (Intel MKL) as the backend support for BLAS and LAPACK related linear algebra operations while for a general purpose with no specific processor types, we rely on OpenBLAS as the backend support for BLAS and LAPACK (via --features “Intel-mkl-static” or “openblas- static”). Intel MKL is slightly faster than OpenBLAS for the testing dataset while OpenBLAS is more flexible and can be used for other aarch64/ARM64 CPUs. Note that both Intel MKL and OpenBLAS implementations take advantage of floating-point hardware such as registers or SIMD instructions. For MacOS, we relied on the accelerate framework in the system as the default backend for BLAS and LAPACK.

### Annembed for metagenomic binning

Metegenomic binning is to cluster similar sequences into bins, which can be thought as a unit for representing population genome sequences. Automatic binners includes MaxBin2, MetaBAT2, CONCOCT et.al. while manual binner includes mmgenome2, BinaRena et.al.. Automatic binners rely on a pre-defined threshold (e.g., a probability threshold or clustering threshold based on contig kmer composition and coverage) (Kang, et al., 2019; Wu, et al., 2015) for calling contig sequences from the same true bin. Mannual binners visualize the kmer composition/profile via embedding them into low-dimensional space (e.g., UMAP, t-SNE), together with the coverage of each contig and then clustering the representation in low dimension space or make decision based on intuition of how close they are in the embedded space. We use the standard PacBio SMRT gut metagenomic sequencing dataset as in (Lamurias, et al., 2022) to compare automatic and manual binners.

## Data and code availability

Annembed package can be found here: https://github.com/jean-pierreBoth/annembed. Scripts for reproducing the main analysis can be found here: https://github.com/jianshu93/annembed_analysis.

## Author Contribution

J.Z, J. P-B and K.T.K designed the work. J.P-B and J.Z wrote the code (genomics par and algorithm part respectively). J.Z did the analysis and benchmark. J.Z and K.T.K wrote the manuscript.

## Acknowledgment

We want to thank PACE at Georgia Tech for providing computational resources. This work was supported by US National Science Foundation (Award No 1759831 and 2129823 to KTK)

---

**Table 1.** Running time of UMAP and annembed for 3 benchmark datasets using 24 threads. Running time is average values of 3 independent runs.

| Dataset | Number of data points | dimension | K | UMAP (NN-Descent) NNS<sup>a</sup> | UMAP (annoy) NNS | UMAP embedding<sup>b</sup> | Annembed (HNSW) NNS | Annembed (embedding) |
| --- | --- | --- | --- | --- | --- | --- | --- | --- |
| MNIST<br>FASHION | 70,000 | 748 | 15 | 2 min 25 s | 44 s | 1 min 01s | 27.4s | 14.2 s |
| SIFT_1M | 1 M | 128 | 15 | 53 min 20 s | 14 min 35s | 53 min 22s | 9 min 21s | 16 min 13s |
| HIGGS | 11 M | 20 | 15 | > 8 h | 1 h 45 min | > 24 h | 43 min | 1 h 56 min |

<sup>a</sup>NN-descent algorithm is not parallelized, and it can only use 1 thread. <sup>b</sup>embedding step in both python version UMAP and R version UWOT is not parallelized.

## Bibliography

1. Amid, E. and Warmuth, M.K. TriMap: Large-scale dimensionality reduction using triplets. arXiv preprint arXiv:1910.00204 2019.

2. Amsaleg, L., et al. Estimating local intrinsic dimensionality. In, Proceedings of the 21th ACM SIGKDD International Conference on Knowledge Discovery and Data Mining. 2015. p. 29–38.

3. Amsaleg, L., et al. Intrinsic dimensionality estimation within tight localities. In, *Proceedings of the 2019 SIAM international conference on data mining*. SIAM; 2019. p. 181–189.

4. Argerich, L. and Golmar, N. Generic LSH Families for the Angular Distance Based on Johnson-Lindenstrauss Projections and Feature Hashing LSH. arXiv preprint arXiv:1704.04684 2017.

5. Aumüller, M., Bernhardsson, E. and Faithfull, A. ANN-Benchmarks: A benchmarking tool for approximate nearest neighbor algorithms. Information Systems 2020;87:101374.

6. Aumüller, M. and Ceccarello, M. The role of local dimensionality measures in benchmarking nearest neighbor search. Information Systems 2021;101:101807.

7. Becht, E., et al. Dimensionality reduction for visualizing single-cell data using UMAP. Nature biotechnology 2019;37(1):38–44.

8. Böhm, J.N., Berens, P. and Kobak, D. Attraction-repulsion spectrum in neighbor embeddings. The Journal of Machine Learning Research 2022;23(1):4118–4149.

9. Bratić, B., et al. NN-Descent on high-dimensional data. In, Proceedings of the 8th International Conference on Web Intelligence, Mining and Semantics. 2018. p. 1-8.

10. Camargo, A.P., et al. IMG/VR v4: an expanded database of uncultivated virus genomes within a framework of extensive functional, taxonomic, and ecological metadata. Nucleic Acids Research 2022;51(D1):D733–D743.

11. Camastra, F. and Staiano, A. Intrinsic dimension estimation: Advances and open problems. Information Sciences 2016;328:26–41.

12. Chen, J., Fang, H.-r. and Saad, Y. Fast Approximate kNN Graph Construction for High Dimensional Data via Recursive Lanczos Bisection. Journal of Machine Learning Research 2009;10(9).

13. Christiani, T. DartMinHash: Fast Sketching for Weighted Sets. arXiv preprint arXiv:2005.11547 2020.

14. Coifman, R.R., et al. Geometric diffusions as a tool for harmonic analysis and structure definition of data: Diffusion maps. Proceedings of the national academy of sciences 2005;102(21):7426-7431.

15. Damrich, S. and Hamprecht, F.A. On UMAP’s true loss function. Advances in Neural Information Processing Systems 2021;34:5798–5809.

16. Datar, M., et al. Locality-sensitive hashing scheme based on p-stable distributions. In, Proceedings of the twentieth annual symposium on Computational geometry. 2004. p. 253–262.

17. Deng, J., et al. Imagenet: A large-scale hierarchical image database. In, 2009 IEEE conference on computer vision and pattern recognition. Ieee; 2009. p. 248-255.

18. Dong, W., Moses, C. and Li, K. Efficient k-nearest neighbor graph construction for generic similarity measures. In, Proceedings of the 20th international conference on World wide web. 2011. p. 577- 586.

19. Dong, W., Moses, C. and Li, K. Efficient k-nearest neighbor graph construction for generic similarity measures. Proceedings of the 20th international conference on World wide web 2011:577-586.

20. Edgar, R. Taxonomy annotation and guide tree errors in 16S rRNA databases. PeerJ 2018;6:e5030.

21. Edgar, R.C. SINTAX: a simple non-Bayesian taxonomy classifier for 16S and ITS sequences. biorxiv 2016:074161.

22. Ertl, O. Superminhash-A new minwise hashing algorithm for jaccard similarity estimation. arXiv preprint arXiv:1706.05698 2017.

23. Ertl, O. BagMinHash - Minwise Hashing Algorithm for Weighted Sets. In, Proceedings of the 24th ACM SIGKDD International Conference on Knowledge Discovery & Data Mining. London, United Kingdom: Association for Computing Machinery; 2018. p. 1368–1377.

24. Ertl, O. ProbMinHash – A Class of Locality-Sensitive Hash Algorithms for the (Probability) Jaccard Similarity. IEEE Transactions on Knowledge and Data Engineering 2020:1–1.

25. Ertl, O. SetSketch: filling the gap between MinHash and HyperLogLog. Proc. VLDB Endow. 2021;14(11):2244–2257.

26. Hajebi, K., et al. Fast approximate nearest-neighbor search with k-nearest neighbor graph. In, Twenty-Second International Joint Conference on Artificial Intelligence. 2011.

27. Halko, N., Martinsson, P.-G. and Tropp, J.A. Finding structure with randomness: Probabilistic algorithms for constructing approximate matrix decompositions. SIAM review 2011;53(2):217–288.

28. Ioffe, S. Improved Consistent Sampling, Weighted Minhash and L1 Sketching. In, 2010 IEEE International Conference on Data Mining. 2010. p. 246-255.

29. Jain, C., et al. High throughput ANI analysis of 90K prokaryotic genomes reveals clear species boundaries. Nature Communications 2018;9(1):5114.

30. Kang, D.D., et al. MetaBAT 2: an adaptive binning algorithm for robust and efficient genome reconstruction from metagenome assemblies. PeerJ 2019;7:e7359.

31. Karst, S.M., Kirkegaard, R.H. and Albertsen, M. Mmgenome: a toolbox for reproducible genome extraction from metagenomes. BioRxiv 2016:059121.

32. Kobak, D. and Berens, P. The art of using t-SNE for single-cell transcriptomics. Nature communications 2019;10(1):1–14.

33. Koonin, E.V. and Wolf, Y.I. Genomics of bacteria and archaea: the emerging dynamic view of the prokaryotic world. Nucleic Acids Research 2008;36(21):6688–6719.

34. Koslicki, D. and Zabeti, H. Improving MinHash via the containment index with applications to metagenomic analysis. Applied Mathematics and Computation 2019;354:206–215.

35. Lamurias, A., et al. Metagenomic binning with assembly graph embeddings. Bioinformatics 2022;38(19):4481–4487.

36. Levina, E. and Bickel, P. Maximum likelihood estimation of intrinsic dimension. Advances in neural information processing systems 2004;17.

37. Li, X. and Li, P. C-MinHash: Improving Minwise Hashing with Circulant Permutation. In, International Conference on Machine Learning. PMLR; 2022. p. 12857-12887.

38. Lin, P.-C. and Zhao, W.-L. Graph based nearest neighbor search: Promises and failures. arXiv preprint arXiv:1904.02077 2019.

39. Malkov, Y.A. and Yashunin, D.A. Efficient and Robust Approximate Nearest Neighbor Search Using Hierarchical Navigable Small World Graphs. IEEE Transactions on Pattern Analysis and Machine Intelligence 2020;42(4):824–836.

40. Marçais, G., et al. Locality-sensitive hashing for the edit distance. Bioinformatics 2019;35(14):i127–i135.

41. McInnes, L., Healy, J. and Melville, J. Umap: Uniform manifold approximation and projection for dimension reduction. arXiv preprint arXiv:1802.03426 2018.

42. Moulton, R. and Jiang, Y. Maximally Consistent Sampling and the Jaccard Index of Probability Distributions. In, 2018 IEEE International Conference on Data Mining (ICDM). 2018. p. 347-356. Murray, C.S., Gao, Y. and Wu, M. Re-evaluating the evidence for a universal genetic boundary among microbial species. Nature Communications 2021;12(1):4059.

43. Newell, R.J.P., Tyson, G. W., & Woodcroft, B. J. Rosella: Metagenomic binning using UMAP and HDBSCAN. zendo 2023.

44. Ondov, B.D., et al. Mash: fast genome and metagenome distance estimation using MinHash. Genome Biology 2016;17(1):132.

45. Pacuk, A., et al. Locality-Sensitive Hashing Without False Negatives for L_p. In, International Computing and Combinatorics Conference. Springer; 2016. p. 105-118.

46. Pagh, R. Locality-sensitive hashing without false negatives. In, Proceedings of the twenty-seventh annual ACM-SIAM symposium on Discrete algorithms. SIAM; 2016. p. 1–9.

47. Parks, D.H., et al. GTDB: an ongoing census of bacterial and archaeal diversity through a phylogenetically consistent, rank normalized and complete genome-based taxonomy. Nucleic Acids Research 2021.

48. Pavia, M.J., et al. BinaRena: a dedicated interactive platform for human-guided exploration and binning of metagenomes. Microbiome 2023;11(1):186.

49. Radovanovic, M., Nanopoulos, A. and Ivanovic, M. Hubs in space: Popular nearest neighbors in high-dimensional data. Journal of Machine Learning Research 2010;11(sept):2487–2531.

50. Rowe, W.P.M., et al. Streaming histogram sketching for rapid microbiome analytics. Microbiome 2019;7(1):40.

51. Schmartz, G.P., et al. BusyBee Web: towards comprehensive and differential composition-based metagenomic binning. Nucleic Acids Research 2022.

52. Shrivastava, A. Optimal densification for fast and accurate minwise hashing. International Conference on Machine Learning 2017:3154–3163.

53. Tan, S., et al. Norm Adjusted Proximity Graph for Fast Inner Product Retrieval. In, Proceedings of the 27th ACM SIGKDD Conference on Knowledge Discovery & Data Mining. Virtual Event, Singapore: Association for Computing Machinery; 2021. p. 1552–1560.

54. Tang, J., et al. Visualizing large-scale and high-dimensional data. In, Proceedings of the 25th international conference on world wide web. 2016. p. 287-297.

55. Van der Maaten, L. and Hinton, G. Visualizing data using t-SNE. Journal of machine learning research 2008;9(11).

56. Wang, D., Shi, L. and Cao, J. Fast algorithm for approximate k-nearest neighbor graph construction. In, 2013 IEEE 13th international conference on data mining workshops. IEEE; 2013. p. 349-356.

57. Wang, Q., et al. Naïve Bayesian Classifier for Rapid Assignment of rRNA Sequences into the New Bacterial Taxonomy. Applied and Environmental Microbiology 2007;73(16):5261–5267.

58. Wu, W., et al. A review for weighted minhash algorithms. IEEE Transactions on Knowledge and Data Engineering 2020;34(6):2553–2573.

59. Wu, Y.-W., Simmons, B.A. and Singer, S.W. MaxBin 2.0: an automated binning algorithm to recover genomes from multiple metagenomic datasets. Bioinformatics 2015;32(4):605–607.

60. Yang, D., et al. D2histoSketch: Discriminative and Dynamic Similarity-Preserving Sketching of Streaming Histograms. IEEE Transactions on Knowledge and Data Engineering 2019;31(10):1898–1911.

61. Zhao, J., et al. GSearch: Ultra-Fast and Scalable Microbial Genome Search by combining Kmer Hashing with Hierarchical Navigable Small World Graphs. bioRxiv 2022:2022.2010.2021.513218.

62. Zhao, X., et al. Towards Efficient Index Construction and Approximate Nearest Neighbor Search in High-Dimensional Spaces. VLDB Endowment 2023.

63. Zu, X. and Tao, Q. SpaceMAP: Visualizing high-dimensional data by space expansion. In, Proc. Int. Conf. Mach. Learn.(ICML). 2022. p. 27707–27723.

